# Mutational and Expression Profile of ZNF217, ZNF750, ZNF703 Zinc Finger Genes in Kenyan Women Diagnosed with Breast Cancer

**DOI:** 10.1101/2025.07.17.665450

**Authors:** Michael Kitoi, John Gitau, Godfrey Wagutu, Kennedy Mwangi, Florence Ngonga, Francis Makokha

## Abstract

**Objective:** To characterize the mutational landscape and expression profiles of ZNF217, ZNF703, and ZNF750, and assess their clinical relevance in breast cancer patients from Kenya.

**Methods:** Whole-exome sequencing (WES) and RNA sequencing (RNA-Seq) data from 23 paired tumor–normal samples were analyzed in a Linux-based environment. Somatic mutations were identified using MuTect2 following alignment to the hg38 reference genome and annotation with VEP. Variants were classified by type, coding consequence, and protein position, and mapped to functional domains. Recurrent mutations were identified, and comparisons were made with The Cancer Genome Atlas (TCGA). Gene expression was quantified using STAR and featureCounts, normalized with DESeq2, and analyzed using paired statistical tests with multiple testing correction. Principal component analysis (PCA) and regression analyses were performed to assess expression patterns and clinical associations.

**Results:** ZNF217 and ZNF750 exhibited high mutational burdens, whereas ZNF703 showed a lower mutation frequency. Mutations were predominantly single nucleotide variants, with missense and synonymous variants as the major classes. Variants were distributed across protein sequences, with limited domain enrichment and no clear hotspot clustering. Recurrent mutations were gene-specific and infrequent. Comparison with TCGA data showed concordant mutation prevalence for ZNF217, low frequency for ZNF703, and absence of ZNF750 mutations. All three genes were significantly upregulated in tumors compared to matched normal tissues (ZNF217: p = 0.00068; ZNF703: p = 0.00475; ZNF750: p = 0.00366). Tumor expression exceeded normal expression in 74% of cases for ZNF217, 64% for ZNF703, and 83% for ZNF750. PCA demonstrated partial separation between tumor and normal samples. ZNF703 expression was positively associated with body mass index (β = 0.194, p = 0.025), and ZNF750 expression was higher in estrogen receptor–positive tumors (β = 1.050, p = 0.005).

**Conclusion:** ZNF217, ZNF703, and ZNF750 display distinct mutation and expression profiles in breast cancer, with evidence of cohort-specific variation. These findings highlight gene-specific mechanisms of dysregulation and emphasize the value of integrating genomic and transcriptomic analyses.

## INTRODUCTION

Breast cancer remains a major global health challenge, with approximately 20 million new cancer cases reported in 2022, of which breast cancer accounted for 11.6% and ranked as the second most common malignancy worldwide (Bray et al., 2024). It is also a leading cause of cancer-related mortality, contributing 6.7% of the estimated 9.7 million deaths globally. The burden is particularly pronounced in low- and middle-income countries, including Kenya, where breast cancer is the most frequently diagnosed malignancy among women, accounting for 25.6% of new cases and a substantial proportion of cancer-related deaths (Global Cancer Observatory, 2022). Despite this high burden, the molecular drivers of breast cancer in African populations remain underexplored.

Advances in cancer genomics have highlighted the critical role of zinc finger (ZNF) proteins in tumorigenesis, particularly through their functions as transcriptional regulators that influence cell proliferation, differentiation, and survival (Stradella et al., 2022). Among these, ZNF217, ZNF703, and ZNF750 have emerged as key genes implicated in breast cancer, yet their precise mutational landscapes and functional roles remain incompletely characterized, especially in African cohorts. ZNF217 is a well-established oncogenic regulator located on chromosome 20q13, a region frequently amplified in breast cancer and associated with poor prognosis (Krig et al., 2010). As a transcriptional co-regulator, ZNF217 interacts with chromatin-modifying complexes to repress or activate gene expression programs involved in cell proliferation, apoptosis resistance, and genomic stability (Krig et al., 2010; Ansari-Pour et al., 2021). Amplification and overexpression of ZNF217 have been shown to promote oncogenic signaling pathways, including activation of ERBB3 and PI3K/AKT signaling, thereby enhancing tumor cell survival and metastatic potential (Krig et al., 2010). In addition to copy number alterations, structural variants such as missense and frameshift mutations can further disrupt its regulatory domains, leading to aberrant transcriptional control and increased tumor aggressiveness (Ansari-Pour et al., 2021). Notably, ZNF217 alterations have been reported to be more prevalent in tumors from individuals of African ancestry, where they have been linked to more aggressive disease phenotypes and poorer clinical outcomes (Ansari-Pour et al., 2021).

In contrast, ZNF750, located on chromosome 17q25.3, functions predominantly as a tumor suppressor by regulating epithelial differentiation and maintaining cellular homeostasis (Butera et al., 2020). It plays a critical role in promoting terminal differentiation through transcriptional activation of differentiation-associated genes and repression of progenitor cell programs (Cassandri et al., 2020). Loss-of-function mutations in ZNF750, including truncating and missense variants, have been implicated in impaired differentiation and increased proliferative capacity, particularly in aggressive breast cancer subtypes such as triple-negative breast cancer (Butera et al., 2020; Cassandri et al., 2020). Meanwhile, ZNF703, located on chromosome 8p12, acts as an oncogenic transcription factor and is frequently amplified in luminal B breast cancer (Klæstad et al., 2021). Its overexpression has been shown to drive tumorigenesis by repressing differentiation pathways and activating proliferation-related gene networks, thereby enhancing cell cycle progression and survival (Zhang et al., 2022). ZNF703 amplification is also associated with endocrine resistance and poor clinical outcomes, highlighting its role as a key driver of aggressive tumor behavior within specific molecular subtypes of breast cancer (Klæstad et al., 2021; Zhang et al., 2022).

Although these genes have been studied individually, several critical gaps remain. First, there is limited understanding of the comprehensive mutation spectrum, including substitution patterns and protein-domain localization, of these genes in African populations. Secondly, the relationship between mutational profiles and gene expression dysregulation still remains unclear. Third, existing studies have largely focused on Western populations, leaving a gap in knowledge regarding population-specific genomic variation and its implications for disease biology and clinical outcomes in sub-Saharan Africa. Addressing these gaps is essential for improving the understanding of breast cancer biology in underrepresented populations and for advancing precision oncology approaches. Therefore, this study aimed to characterize the somatic mutation landscape of ZNF217, ZNF703, and ZNF750, including mutation types, frequencies, and distribution across protein domains; evaluate their expression patterns in tumor and matched normal tissues; and assess associations between gene alterations and clinicopathological variables in a cohort of Kenyan breast cancer patients.

## MATERIALS AND METHODS

### Patients and Samples

This research collected its data from a pilot study examining breast cancer progression in Kenyan patients between 2019 and 2021 under informed consent and the study was approved by Research and Ethics Committees at Aga Khan University Hospital, Nairobi (Ref: 2018/REC-80) and AIC Kijabe Hospital (KH IERC-02718/2019). After patients granted their consent at AIC Kijabe Hospital and Aga Khan University Hospital, Nairobi, purposive sampling enrolled 23 patients with malignant breast cancer. Fresh tumor tissue together with adjacent normal tissue were extracted from 23 patients, while 46 patient samples were sent to the National Cancer Institute at Bethesda MD USA for sequencing procedures.

### Whole-exome sequencing and RNA-sequencing

Whole-exome sequencing (WES) and RNA sequencing (RNA-seq) were performed on paired tumor and adjacent normal tissue samples from 23 patients. Genomic DNA was extracted from tissue samples using the DNeasy Blood and Tissue Kit (Qiagen, Hilden, Germany) according to the manufacturer’s instructions. Total RNA was extracted from frozen tissue samples using TRIzol reagent (Invitrogen), following standard protocols. Whole-exome sequencing (WES) was performed by Psomagen Inc. (https://www.psomagen.com/). Library preparation, target enrichment, and sequencing were conducted using established protocols consistent with high-throughput Illumina sequencing workflows as was previously described by Tang et al. (2023). Sequencing generated paired-end reads of 150 base pairs. Tumor samples were sequenced at an average depth of approximately 250×, while matched normal samples were sequenced at approximately 150× coverage.

For RNA samples, quality and integrity were assessed using the Agilent 2100 Bioanalyzer, and only samples with RNA integrity number (RIN) ≥ 7 were included for sequencing. RNA-seq libraries were prepared using the TruSeq PolyA kit (Illumina) mRNA Library Preparation Kit, which includes poly-A selection, RNA fragmentation, first- and second-strand cDNA synthesis, end repair, adapter ligation, and PCR amplification. Sequencing was performed on the Illumina NovaSeq platform, generating paired-end reads of 150 base pairs. Tumor samples were sequenced at an average depth of approximately 250× for WES and ∼30 million reads for RNA-seq, while normal samples were sequenced at ∼150× depth for WES.

The sequencing raw data and associated sample descriptors used in this study were previously generated by Tang et al. (2023) and are publicly available in the Sequence Read Archive (SRA, RRID:SCR_004891) at the National Center for Biotechnology Information (NCBI) under accession number PRJNA913947.

### Mutation Analysis

All analyses were performed in a Linux-based computational environment. Raw sequencing reads were quality-checked using FastQC (v0.11.9) and summarized with MultiQC (v1.14). Adapter sequences and low-quality bases were trimmed using Trimmomatic (v0.39) with parameters: leading/trailing quality <3, sliding window (4:15), and minimum length of 36 bp. Trimmed reads were aligned to the human reference genome (hg38) using BWA-MEM (v0.7.17). Alignments were processed using SAMtools (v1.15) for sorting and indexing, and duplicate reads were marked with Picard (v2.26.10). Base quality score recalibration was performed using GATK (v4.2) following best practices.

Somatic variants were called using MuTect2 in paired tumor–normal mode. A panel of normals (PON) was generated to remove recurrent artifacts and germline variants. Variants were filtered using GATK FilterMutectCalls with thresholds of minimum base quality 20, read depth ≥10, and variant allele fraction ≥0.05. Variants were normalized using VT and annotated with Ensembl VEP (v108). All variants were visually inspected using the Integrative Genomics Viewer (IGV) to confirm read support and alignment quality.

For downstream analyses, variants were classified as SNPs or indels, and coding consequences were grouped into synonymous, missense, frameshift, and in-frame deletion categories. Mutation burden per gene and per patient was calculated, and nucleotide substitution patterns were summarized to generate mutation spectra. Amino acid positions were extracted from annotated protein changes and mapped onto protein sequences of ZNF217, ZNF703, and ZNF750. Functional domain annotations were obtained from UniProt, and mutations were classified as occurring within or outside domains.

Recurrent mutations were identified by aggregating identical amino acid changes across samples. TCGA mutation data were curated and standardized to enable comparison of gene-level mutation frequency, mutation types, and protein-level distribution. All statistical analyses and visualizations were performed in R (v4.x) using ggplot2, dplyr, and tidyr.

### Gene Expression Analysis

All gene expression analyses were conducted in R statistical software (version 4.5.0). RNA sequencing reads were aligned to the human reference genome (hg38) using the STAR aligner (v2.7.10a), which performs spliced alignment and generates alignment metrics. Gene-level quantification was performed using featureCounts (v2.0.3) with GENCODE gene annotations to assign reads to genomic features and generate raw count data. Raw count data were imported into R (v4.5.0) and normalized using the DESeq2 package (v1.40), which applies the median-of-ratios method to correct for differences in sequencing depth and library size. The resulting normalized count matrix was used for downstream analyses.

Differential gene expression analysis was conducted using paired tumor–normal comparisons. The normality of paired differences was assessed using the Shapiro–Wilk test. Where normality assumptions were met, paired t-tests were applied; otherwise, the Wilcoxon signed-rank test was used. P-values were adjusted for multiple comparisons using the Benjamini–Hochberg false discovery rate (FDR) method, and an adjusted p-value of ≤ 0.05 was considered statistically significant. Data visualization, including boxplots, violin plots, heatmaps, and principal component analysis (PCA), was performed using the ggplot2 and pheatmap packages in R.

## RESULTS

### Patient Demographic and Clinical Characteristics

A total of 23 women diagnosed with breast cancer were included in this study. The summary statistics of continuous variables are presented in Table 1. The mean age of the cohort was 48.7 years (range: 30–72 years), with a median age of 45 years. The mean body mass index (BMI) was 27.8 kg/m² (range: 21.1–37.8 kg/m²), indicating that a substantial proportion of patients were above normal weight.

**Table 1:** Summary Statistics of Continuous Variables.

| Variable | Mean | Median | Minimum | Maximum | Standard Deviation |
| --- | --- | --- | --- | --- | --- |
| Age (years) | 48.7 | 45 | 30 | 72 | 10.8 |
| BMI (kg/m<sup>2</sup>) | 27.8 | 27.7 | 21.1 | 37.8 | 4.4 |

The distribution of clinical and pathological characteristics is summarized in Table 2. The largest proportion of patients was aged 40–49 years (30.4%), while patients younger than 40 years and those aged ≥60 years each accounted for 21.7% of the cohort. BMI classification showed that 39.1% of patients were overweight and 26.1% were obese, while 34.8% had normal BMI. Tumor grading revealed that the majority of patients (65.2%) presented with Grade 3 tumors, while 34.8% had Grade 2 tumors. In terms of cancer stage, most patients were diagnosed at Stage II (47.8%), followed by Stage III (30.4%) and Stage I (21.7%).

**Table 2:** Summary Distribution of Clinical and Pathological Characteristics.

| Variable | Category | Frequency (n) | Percentage (%) |
| --- | --- | --- | --- |
| Age Group (years) | < 40 | 5 | 21.7 |
|  | 40–49 | 7 | 30.4 |
|  | 50–59 | 6 | 26.1 |
|  | ≥ 60 | 5 | 21.7 |
| BMI Category | Normal (18.5–24.9) | 8 | 34.8 |
|  | Overweight (25–29.9) | 9 | 39.1 |
|  | Obese (≥30) | 6 | 26.1 |
| Tumor Grade | Grade 2 | 8 | 34.8 |
|  | Grade 3 | 15 | 65.2 |
| Cancer Stage | Stage I | 5 | 21.7 |
|  | Stage II | 11 | 47.8 |
|  | Stage III | 7 | 30.4 |
| HER2 Status | Positive | 12 | 52.2 |
|  | Negative | 11 | 47.8 |
| PR Status | Positive | 16 | 69.6 |
|  | Negative | 7 | 30.4 |
| ER Status | Positive | 12 | 52.2 |
|  | Negative | 11 | 47.8 |

Receptor status distribution is also presented in Table 2, showing that 52.2% of patients were HER2-positive and ER-positive, while PR positivity was observed in 69.6% of patients. Combined receptor subtype classification is shown in Table 3, where 34.8% of patients were triple positive (ER+, PR+, HER2+), and 21.7% were triple negative (ER−, PR−, HER2−).

**Table 3:** Combined Receptor Subtypes.

| Subtype | Frequency (n) | Percentage (%) |
| --- | --- | --- |
| Triple Positive (ER+, PR+, HER2+) | 8 | 34.8 |
| Triple Negative (ER–, PR–, HER2–) | 5 | 21.7 |
| Other Combinations | 10 | 43.5 |

### Mutation burden, prevalence, and spectrum

We first assessed mutation burden across ZNF217, ZNF703, and ZNF750. ZNF217 showed the highest number of mutations (∼195), followed by ZNF750 (∼160), whereas ZNF703 exhibited a markedly lower burden (∼25) (Figure 1A; Table 4, Table S1). Across all genes, single nucleotide polymorphisms (SNPs) predominated, with fewer insertions and deletions (indels). Coding consequence analysis revealed that missense and synonymous mutations were the dominant classes (Figure 1B). Furthermore, analysis of the nucleotide substitution spectrum revealed distinct patterns across genes (Figure 2). In ZNF217, C>T transitions were most frequent (43), followed by C>A (22) and G>A (18), with A>T and T>G absent. ZNF750 showed predominant A>G (38) and T>C (36) substitutions, while several classes (e.g., G>C, T>A) were not detected. In contrast, ZNF703 exhibited fewer substitutions overall, with no dominant class; the most frequent were C>T (6) and T>C (4).

**Figure 1:**
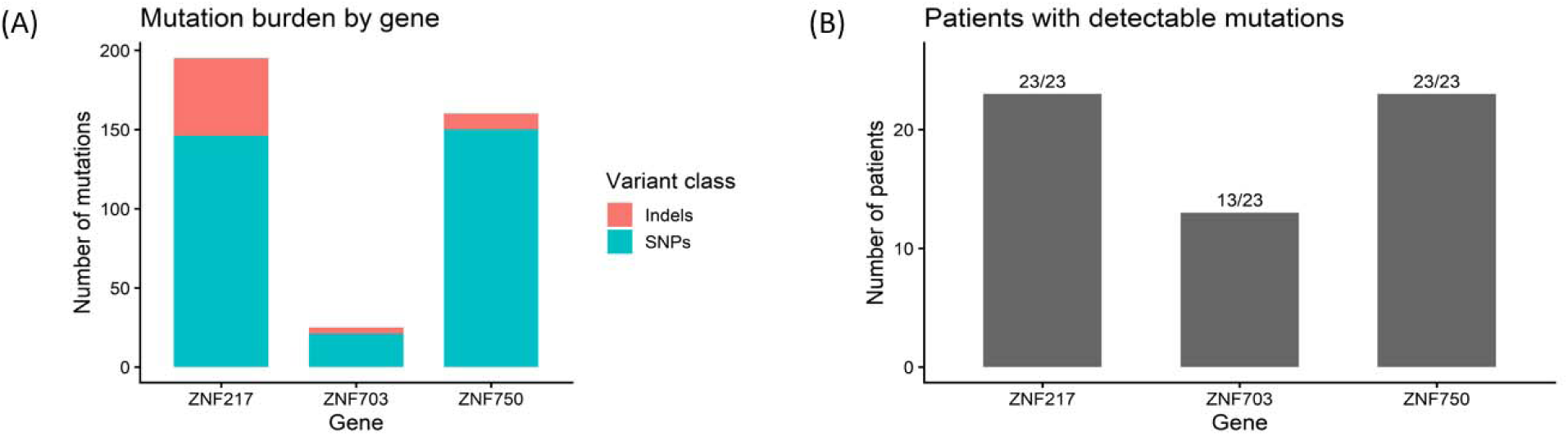
Mutation profile of ZNF217, ZNF703, and ZNF750 genes in 23 patients. (A) Mutation burden by gene showing the number of single nucleotide polymorphisms (SNPs) and indels, (B) Coding consequence composition illustrating the distribution of synonymous, missense, and frameshift variants, (C) Number of patients with detectable mutations per gene.

**Figure 2:**
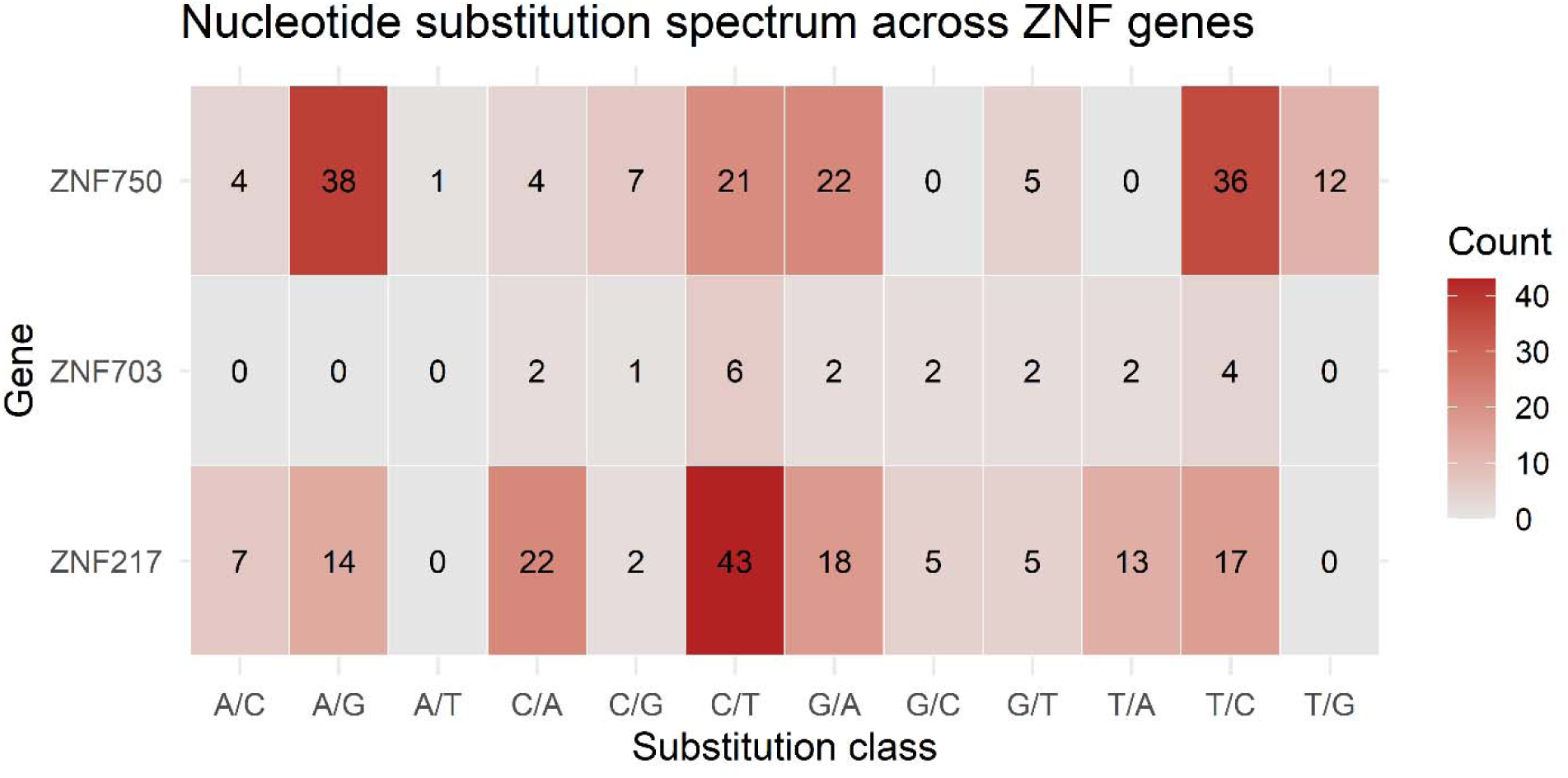
Nucleotide substitution spectrum across ZNF217, ZNF703, and ZNF750 genes in 23 patients. Heatmap showing the distribution and frequency of nucleotide substitution classes (A/C, A/G, A/T, C/A, C/G, C/T, G/A, G/C, G/T, T/A, T/C, T/G) across the three genes, with color intensity representing mutation counts.

**Table 4:** Gene Level Mutation Summary.

| Gene | Total Mutations | Total SNPs | Total Indels | Mutated Patients | Total Patients | Synonymous | Missense | Frameshift | Recurrent Coding Variants | Dominant Substitution |
| --- | --- | --- | --- | --- | --- | --- | --- | --- | --- | --- |
| ZNF217 | 195 | 146 | 49 | 23 | 23 | 13 | 22 | 1 | 3 | C/T |
| ZNF703 | 25 | 21 | 4 | 13 | 23 | 3 | 3 | 1 | 1 | C/T |
| ZNF750 | 160 | 150 | 10 | 23 | 23 | 36 | 17 | 0 | 6 | A/G |

**Table 5:** Recurrent Coding Variants.

| Gene | Mutated Patients | Total Patients | Mutation Frequency | Dataset | Mutation Type | Count |
| --- | --- | --- | --- | --- | --- | --- |
| ZNF217 | 15 | 23 | 0.652174 | This study | Missense | 17 |
| ZNF217 | 15 | 23 | 0.652174 | This study | Synonymous | 4 |
| ZNF750 | 16 | 23 | 0.695652 | This study | Missense | 11 |
| ZNF750 | 16 | 23 | 0.695652 | This study | Synonymous | 31 |
| ZNF217 | 7 | 994 | 0.007042 | TCGA | Missense | 6 |
| ZNF217 | 7 | 994 | 0.007042 | TCGA | Nonsense | 1 |
| ZNF703 | 1 | 994 | 0.001006 | TCGA | Missense | 2 |

### Protein domain distribution and functional enrichment

Mapping mutations onto protein sequences showed that variants were distributed across both domain and non-domain regions (Figure 3, Table S2). In ZNF217, mutations spanned the full protein length, including zinc finger domains, and comprised mainly missense and synonymous variants with a single frameshift mutation. ZNF703 mutations were sparse and scattered, with no clear clustering; most were missense or synonymous, with one in-frame deletion in the C-terminal domain. ZNF750 mutations were distributed throughout the protein, with a subset localized within or near the zinc finger domain. Furthermore, consistent with this, most mutations across all genes occurred outside annotated domains, with only a subset overlapping domain regions (Figure 4A, Table S5). ZNF217 showed limited domain-associated mutations, whereas none were observed for ZNF703 and ZNF750. Mutation composition analysis confirmed the predominance of missense and synonymous variants (Figure 4B), with ZNF217 enriched for missense mutations, ZNF750 for synonymous variants, and ZNF703 showing a more balanced distribution with minor frameshift contribution.

**Figure 3:**
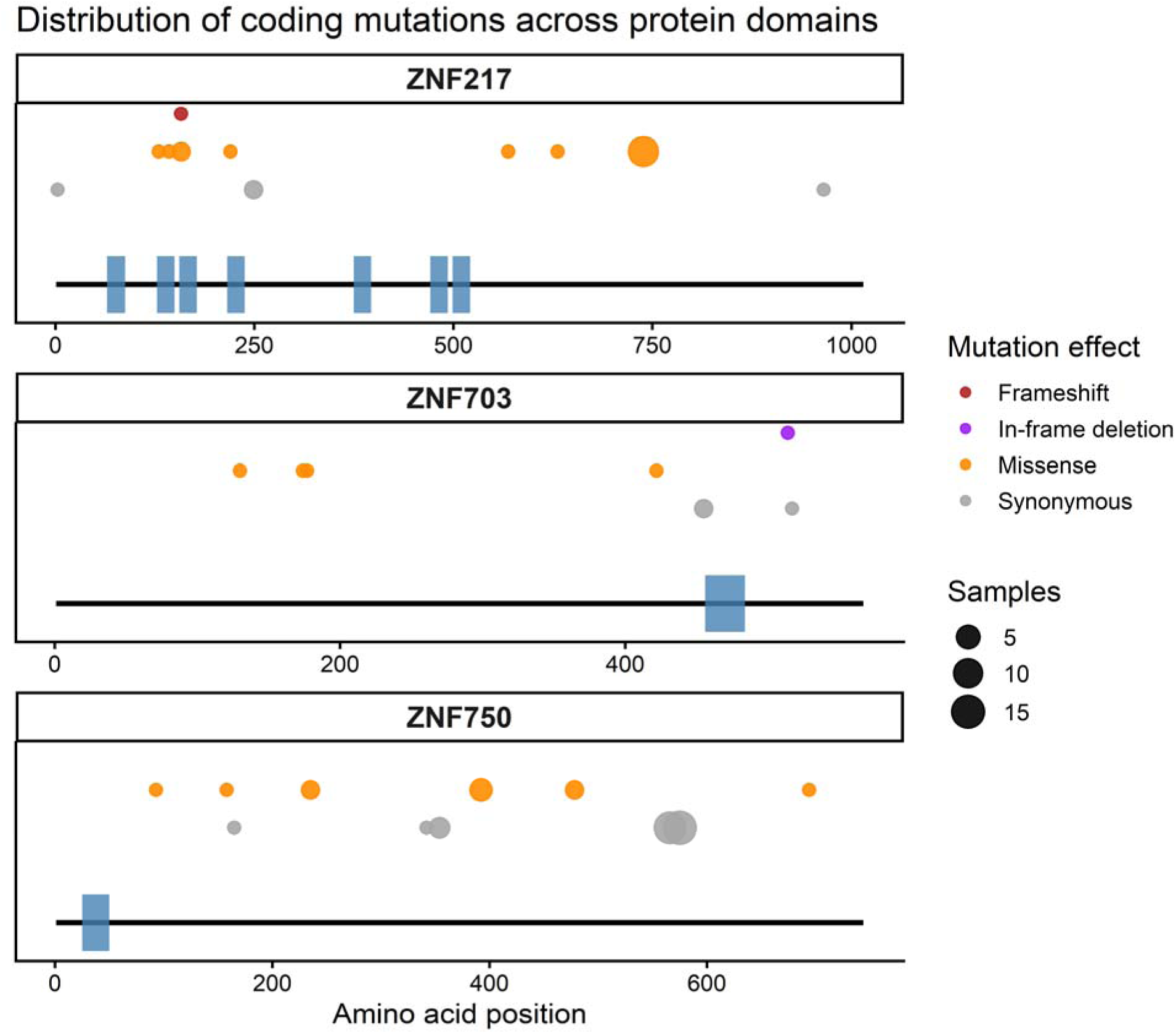
Distribution of coding mutations across protein domains in ZNF217, ZNF703, and ZNF750 genes. Lollipop plots showing the positions of coding mutations along the protein sequences of each gene, with annotated functional domains indicated.

**Figure 4:**
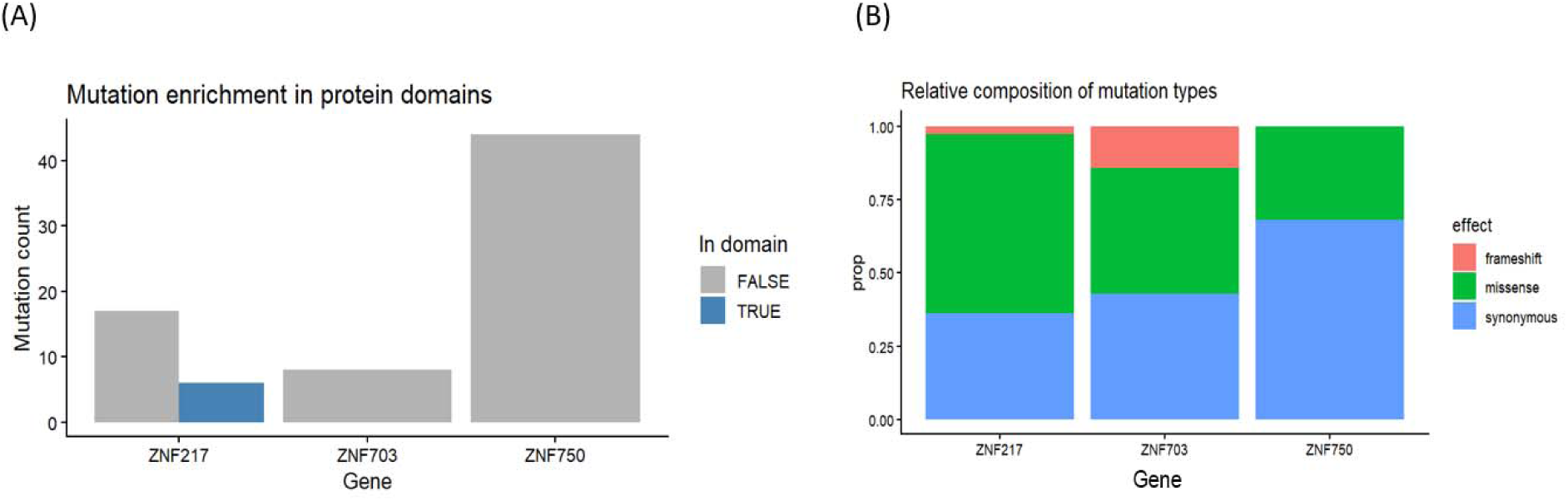
Functional characterization of mutations across ZNF217, ZNF703, and ZNF750 genes. (A) Mutation enrichment in protein domains showing the number of mutations occurring within and outside annotated functional domains for each gene, (B) Relative composition of mutation types illustrating the proportion of synonymous, missense, and frameshift variants across the three genes.

### Recurrent mutations and comparison with TCGA

Recurrent mutation analysis identified several variants present in multiple samples (Figure 5, Table S3). ZNF750 showed the highest recurrence, particularly for synonymous variants such as Ala575= and Pro566=. The most frequent non-synonymous recurrent mutation was Val739>Ile in ZNF217. Additional recurrent missense variants included Gln392>Arg, Val478>Ile, and Met235>Val in ZNF750, whereas recurrence in ZNF703 was minimal, with only a few variants such as Pro455= observed more than once. Comparison with TCGA data demonstrated both concordance and divergence (Table 6, Table S4). ZNF217 remained one of the most frequently altered genes across datasets, while ZNF703 mutations were consistently infrequent. In contrast, although ZNF750 exhibited a high mutation burden and was present in all patients in this cohort, no mutations were detected for ZNF750 in the TCGA dataset, indicating a cohort-specific pattern.

**Figure 5:**
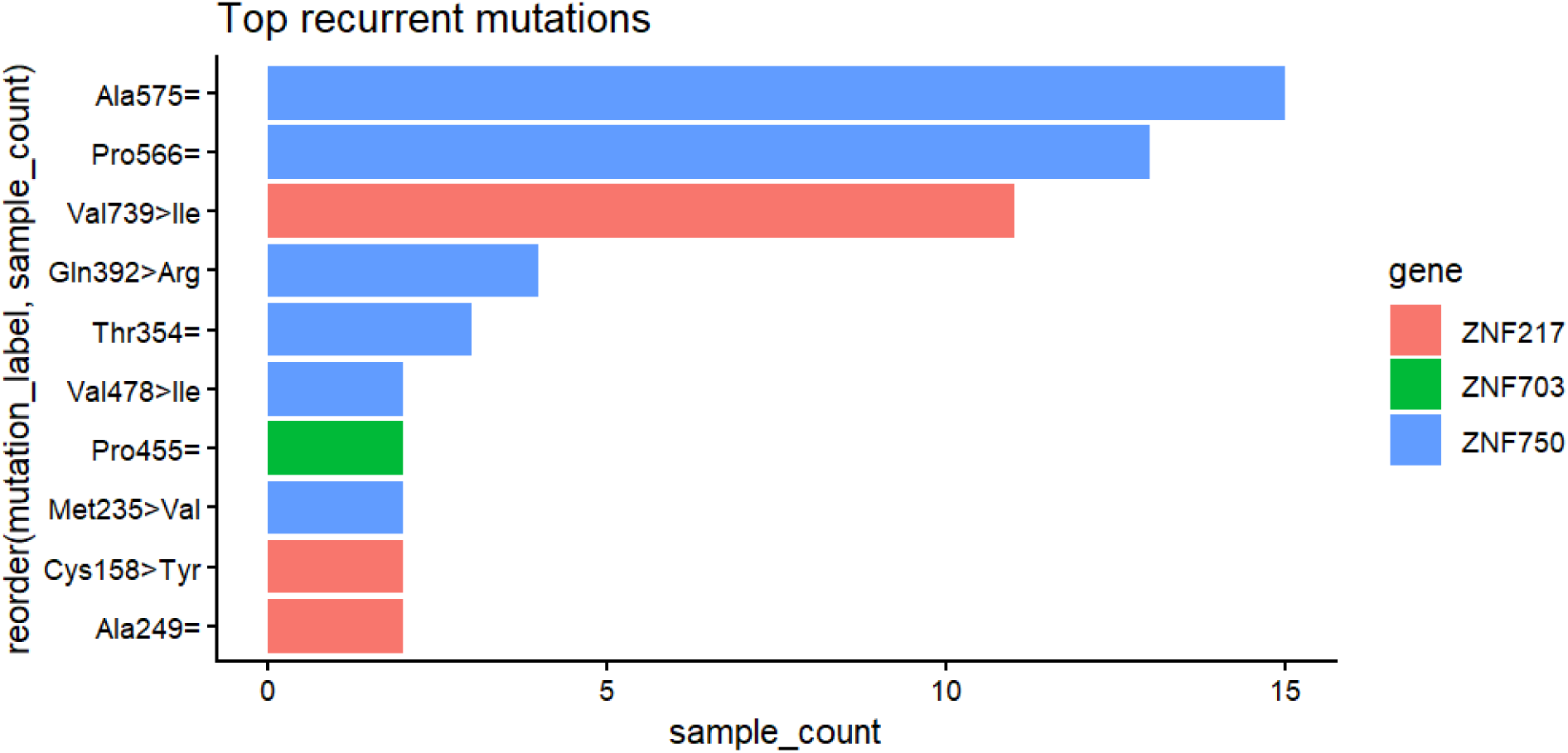
Recurrent mutations across ZNF217, ZNF703, and ZNF750 genes in 23 patients. Bar plot showing the most frequently observed amino acid substitutions across the three genes, with bar length representing the number of samples harboring each mutation and colors indicating gene origin.

**Table 6:** Combined Mutation Comparison.

| Gene | Mutated Patients | Total Patients | Mutation Frequency | Dataset | Mutation Type | Count |
| --- | --- | --- | --- | --- | --- | --- |
| ZNF217 | 15 | 23 | 0.652174 | This study | Missense | 17 |
| ZNF217 | 15 | 23 | 0.652174 | This study | Synonymous | 4 |
| ZNF750 | 16 | 23 | 0.695652 | This study | Missense | 11 |
| ZNF750 | 16 | 23 | 0.695652 | This study | Synonymous | 31 |
| ZNF217 | 7 | 994 | 0.007042 | TCGA | Missense | 6 |
| ZNF217 | 7 | 994 | 0.007042 | TCGA | Nonsense | 1 |
| ZNF703 | 1 | 994 | 0.001006 | TCGA | Missense | 2 |

### Differential Expression of ZNF217, ZNF703, and ZNF750

Differential expression analysis of ZNF217, ZNF703, and ZNF750 was performed using normalized RNA-seq data from 23 paired tumor and matched normal breast tissue samples. All three genes demonstrated significantly higher expression in tumor tissues compared to normal tissues (Table 7). Mean tumor expression values exceeded normal expression for ZNF217 (10.88 vs 10.23), ZNF703 (11.31 vs 10.39), and ZNF750 (7.46 vs 7.02), with positive mean paired differences observed across all genes. At the individual level, tumor expression exceeded matched normal expression in the majority of samples, with 74% of cases for ZNF217, 64% for ZNF703, and 83% for ZNF750 (Table 7). Paired statistical testing confirmed significant differential expression for ZNF217 (p = 0.00068), ZNF703 (p = 0.00475), and ZNF750 (p = 0.00366), with confidence intervals excluding zero (Table 8). These differences remained significant following Benjamini–Hochberg correction (Table 8), indicating robust upregulation of all three genes in tumor samples. Effect size estimates were moderate to large, supporting the magnitude of these expression differences (Table 8). Among the three genes, ZNF703 exhibited the largest magnitude of differential expression, indicating a stronger transcriptional shift in tumor tissues.

**Table 7:** ZNF Expression Descriptive Statistics.

| Gene | n | Mean Tumor | SD Tumor | Mean Normal | SD Normal | Mean Difference | SD Diff | Median Difference | Tumor Higher (%) |
| --- | --- | --- | --- | --- | --- | --- | --- | --- | --- |
| ZNF217 | 23 | 10.8847 | 0.83289 | 10.2301 | 0.2969 | 0.6546 | 0.7948 | 0.4888 | 73.9 |
| ZNF703 | 23 | 11.3147 | 1.5103 | 10.3862 | 0.5878 | 0.9285 | 1.4177 | 0.8957 | 64.3 |
| ZNF750 | 23 | 7.4563 | 0.6622 | 7.0198 | 0.3909 | 0.4365 | 0.6440 | 0.4391 | 82.6 |

**Table 8:** ZNF Expression Descriptive Statistics.

| Gene | p-value | 95% CI (Lower) | 95% CI (Upper) | Mean Difference | SD Difference | Cohen's d | Benjamin–Hochberg |  |
| --- | --- | --- | --- | --- | --- | --- | --- | --- |
|  |  |  |  |  |  |  | Raw p-value | Adjusted p-value |
| ZNF217 | 0.0006 | 0.3109 | 0.9984 | 0.6547 | 0.7949 | 0.8236 | 0.0009 | 0.0029 |
| ZNF703 | 0.0047 | 0.3154 | 1.5416 | 0.9285 | 1.4178 | 0.6549 | 0.0043 | 0.0043 |
| ZNF750 | 0.0036 | 0.1581 | 0.7151 | 0.4366 | 0.6441 | 0.6778 | 0.00345 | 0.0043 |

### Distribution and Variability of Gene Expression Profiles

Visualization of expression distributions revealed consistent shifts toward higher expression in tumor samples across all three genes. Violin and boxplots (Figure 6A) showed broader and right-shifted distributions in tumors, particularly for ZNF217 and ZNF703, indicating increased variability and elevated expression levels compared to normal tissues. Paired expression plots (Figure 6B) demonstrated that most patients exhibited increased tumor expression relative to matched normal samples, with ZNF703 showing the most pronounced and variable increases. Distribution analyses using histograms and density plots (Figure 6C) confirmed that tumor expression values extended into higher ranges not observed in normal samples, particularly for ZNF703, which displayed the widest spread.

**Figure 6.**
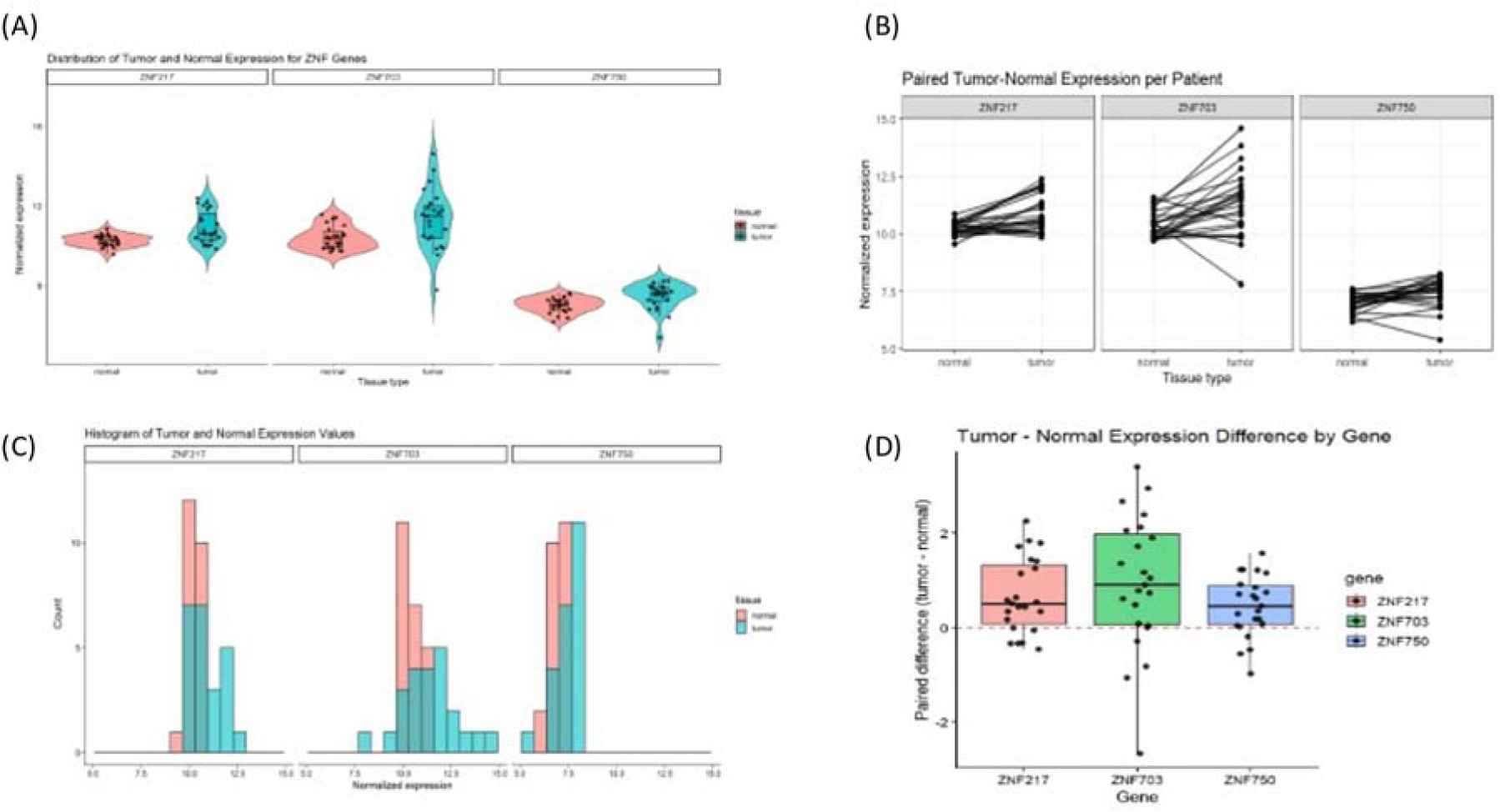
Differential expression patterns of ZNF217, ZNF703, and ZNF750 in paired breast tumor and normal tissues. (A) Violin plots showing the distribution of normalized gene expression levels in tumor and matched normal samples for each gene. (B) Paired tumor–normal expression profiles for individual patients. (C) Histograms depicting the distribution of normalized expression values in tumor and normal tissues. (D) Boxplots of paired expression differences (tumor minus normal) for each gene.

Analysis of paired differences (Figure 6D) further showed that most samples had positive tumor–normal expression differences, with ZNF703 exhibiting the largest variability and highest maximum differences. ZNF750 showed a narrower range of differences, indicating more consistent but smaller expression shifts across patients (Figure 7A). Z-score heatmap analysis (Figure 7B) demonstrated coordinated relative overexpression of all three genes in tumor samples, with variability observed across patients. These results collectively indicate increased expression and heterogeneity of ZNF gene expression in tumor tissues.

**Figure 7.**
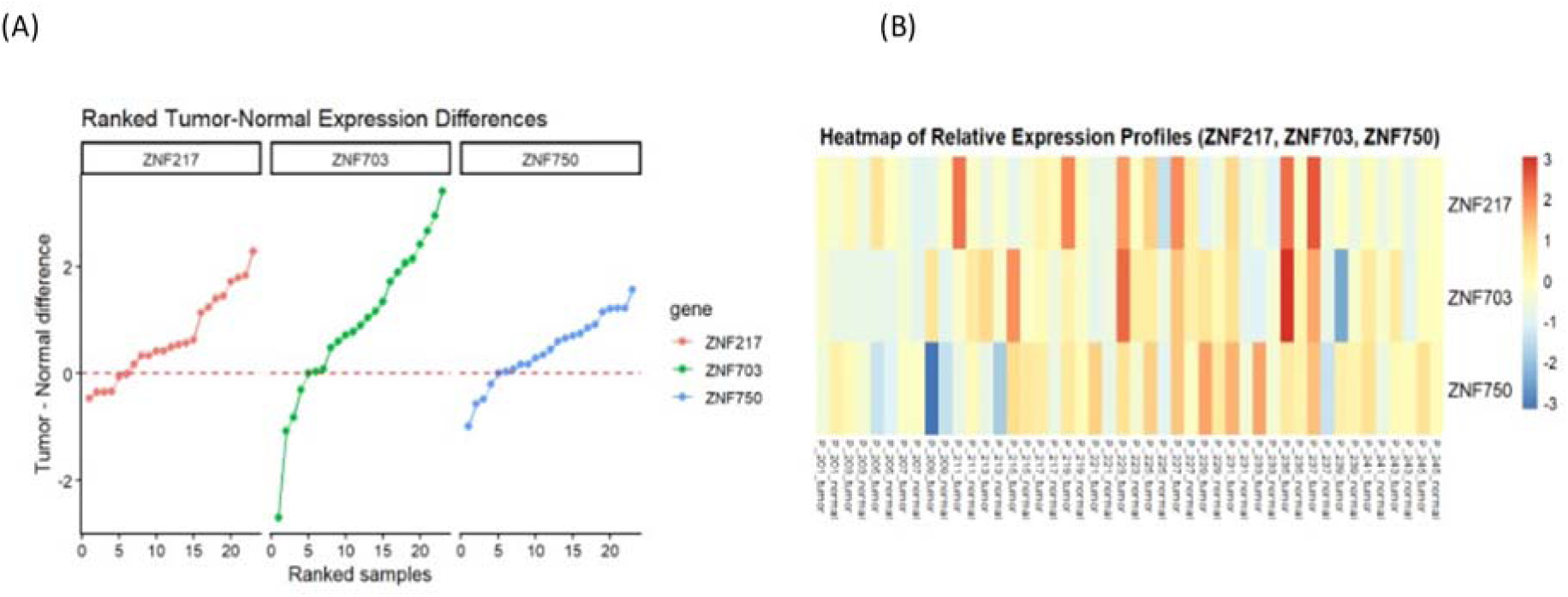
Patient-level expression differences and relative expression patterns of ZNF217, ZNF703, and ZNF750. (A) Ranked tumor–normal expression differences for each gene across individual patients. (B) Heatmap of z-score–normalized expression values for ZNF217, ZNF703, and ZNF750 across all samples, with color scale indicating relative expression levels.

### Principal Component Analysis of Gene Expression Profiles

Principal Component Analysis (PCA) was used to evaluate global expression patterns across samples. The first two principal components explained 75.41% and 13.41% of the variance, respectively (Figure 8). PCA revealed partial separation between tumor and normal samples, with tumor samples tending to cluster distinctly along the principal component axes. Despite this separation, some overlap between tumor and normal samples was observed, indicating heterogeneity in expression profiles within the cohort. These findings suggest that expression patterns of ZNF217, ZNF703, and ZNF750 contribute to distinguishing tumor from normal tissue but do not fully segregate the two groups.

**Figure 8.**
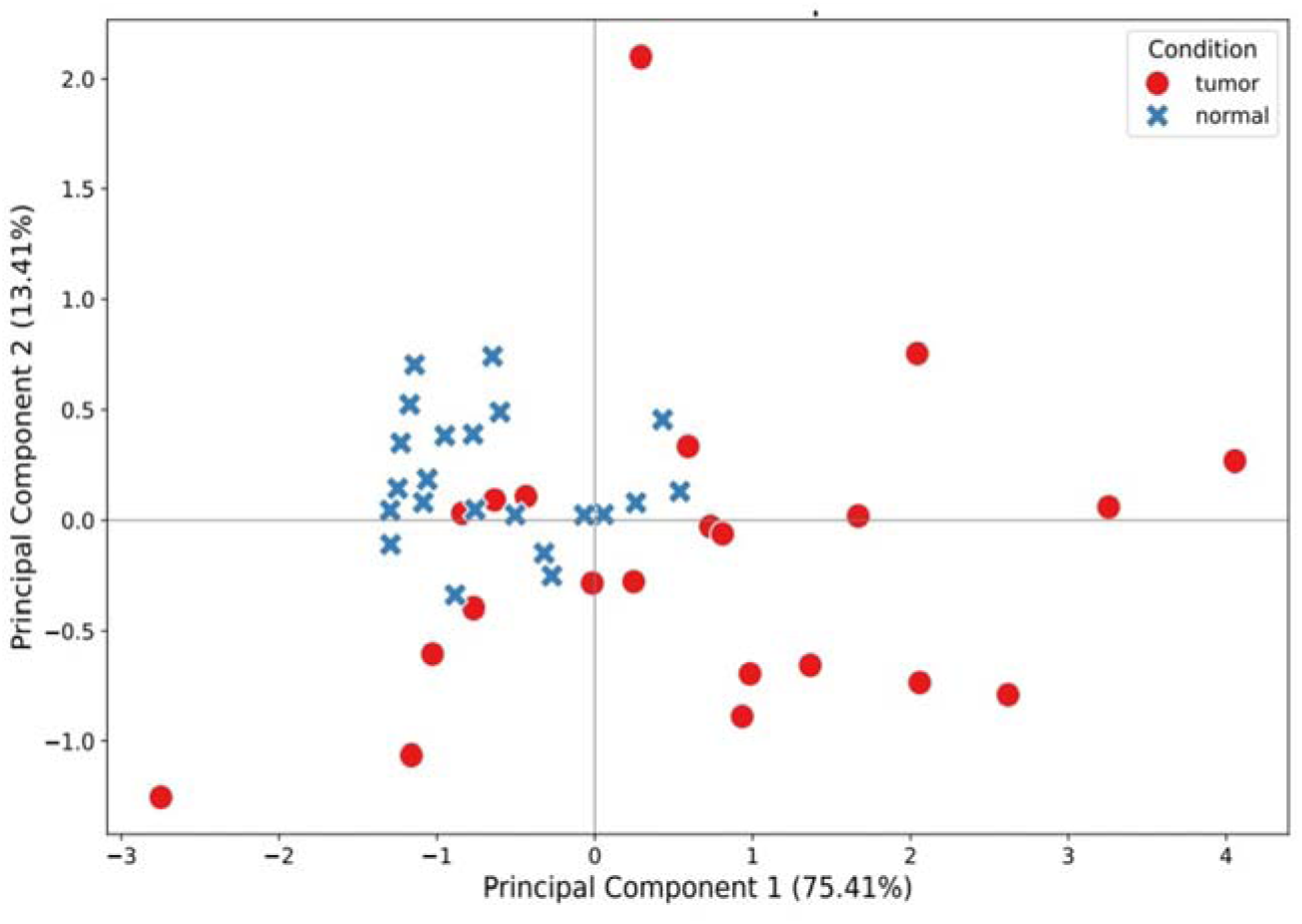
Principal component analysis of gene expression profiles. Scatter plot of the first two principal components based on normalized expression values of ZNF217, ZNF703, and ZNF750, with samples colored by condition (tumor and normal). Axes indicate the proportion of variance explained by each component.

### Clinical Associations of ZNF Gene Expression

Associations between gene expression and clinical variables were generally weak across the cohort. Correlation analysis (Figure 9A) showed a modest positive association between BMI and ZNF703 expression, while ZNF217 and ZNF750 exhibited weaker trends with BMI and age (Figure 9B). Comparisons across tumor grade and cancer stage (Figures 9C and 9D) revealed substantial overlap in expression values across categories, indicating no clear stratification of gene expression by disease severity. ZNF703 consistently showed greater variability across clinical groups, whereas ZNF750 remained more tightly distributed. Similarly, analyses by receptor status (HER2, ER, and PR; Figures 10A–10C) showed overlapping expression distributions across groups for all three genes, with no distinct separation between receptor-positive and receptor-negative tumors. These findings indicate that individual clinical variables do not strongly differentiate expression patterns of the ZNF genes in this cohort.

**Figure 9.**
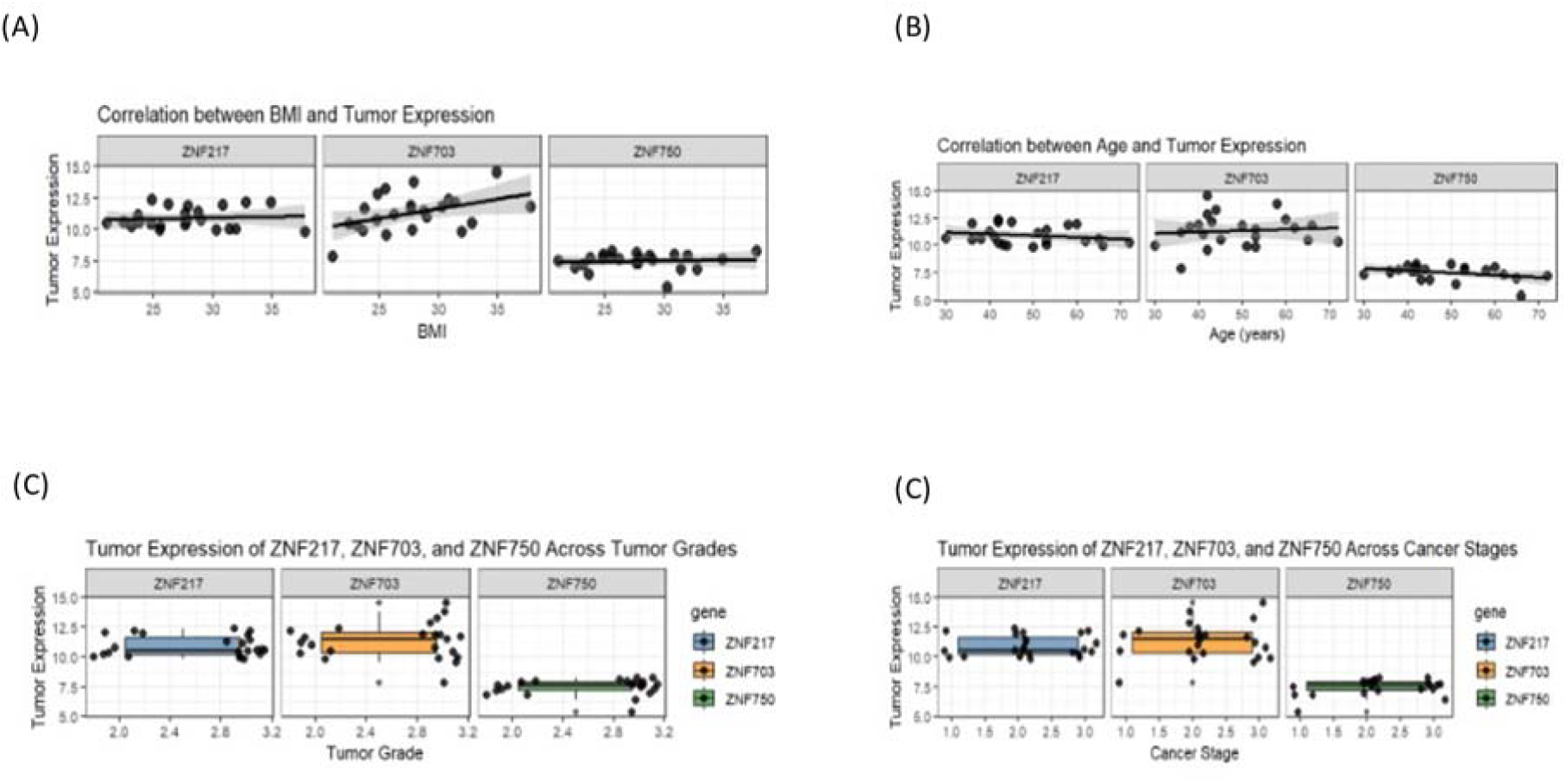
Associations between gene expression and clinical variables. (A) Scatter plots showing the relationship between body mass index (BMI) and tumor expression of ZNF217, ZNF703, and ZNF750. (B) Scatter plots showing the relationship between age and tumor expression for each gene. (C) Boxplots of tumor expression across tumor grades. (D) Boxplots of tumor expression across cancer stages.

**Figure 10.**
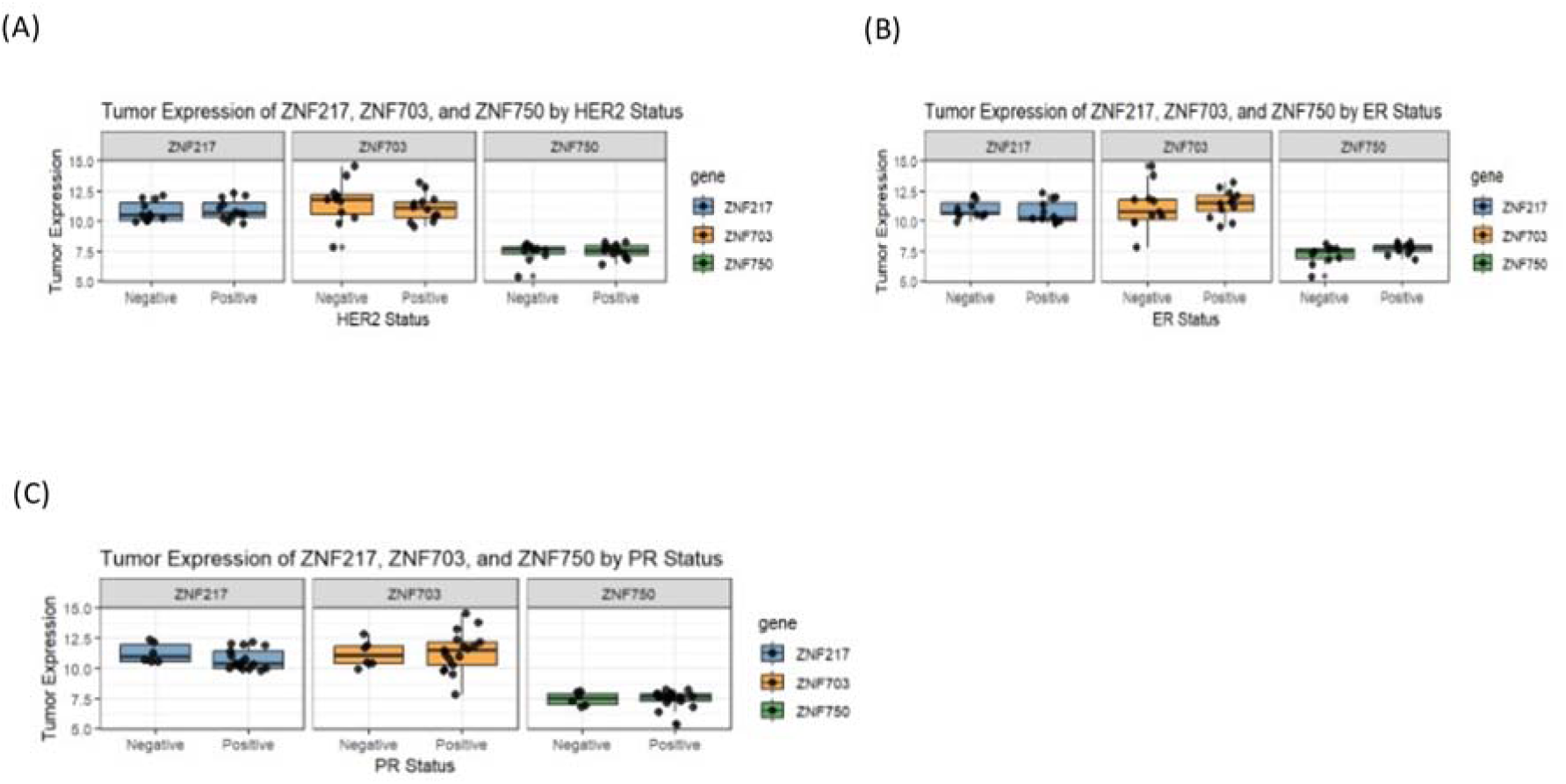
Gene expression across receptor status categories. (A) Boxplots of tumor expression of ZNF217, ZNF703, and ZNF750 stratified by HER2 status. (B) Boxplots of tumor expression stratified by estrogen receptor (ER) status. (C) Boxplots of tumor expression stratified by progesterone receptor (PR) status.

### Multivariable Analysis and Integrated Clinical Associations

Multiple linear regression analysis identified limited independent clinical predictors of gene expression. BMI was significantly associated with increased ZNF703 expression (β = 0.194, p = 0.025), while ER-positive status was significantly associated with higher ZNF750 expression (β = 1.050, p = 0.005) (Table 9, Table S6). No significant predictors were identified for ZNF217. A clinical association heatmap (Figure 11) summarized these relationships, highlighting BMI as the strongest correlate of ZNF703 expression and ER status as the strongest correlate of ZNF750 expression. ZNF217 showed weak correlations with all clinical variables, including a modest negative association with PR status (ρ = −0.36). These findings suggest that while certain clinical variables are associated with gene expression, the overall influence of clinical factors on ZNF gene expression is limited in this cohort.

**Table 9:** Multiple Regression Summary.

| Gene | Variable | Estimate ($\beta$) | Std. Error | t-Statistic | p-value |
| --- | --- | --- | --- | --- | --- |
| **ZNF217** | Intercept | 11.747 | 2.496 | 4.706 | 0.0003 |
|  | Age | -0.009 | 0.019 | -0.478 | 0.640 |
|  | BMI | 0.027 | 0.050 | 0.539 | 0.598 |
|  | Tumor Grade | -0.403 | 0.505 | -0.798 | 0.437 |
|  | Cancer Stage | 0.174 | 0.321 | 0.543 | 0.595 |
|  | HER2 Positive | 0.072 | 0.481 | 0.149 | 0.883 |
|  | ER Positive | -0.357 | 0.551 | -0.647 | 0.528 |
|  | PR Positive | -0.442 | 0.562 | -0.786 | 0.444 |
| **ZNF703** | Intercept | 1.394 | 3.854 | 0.362 | 0.723 |
|  | Age | 0.025 | 0.030 | 0.827 | 0.421 |
|  | BMI | 0.194 | 0.078 | 2.497 | 0.025 |
|  | Tumor Grade | 0.969 | 0.780 | 1.243 | 0.233 |
|  | Cancer Stage | 0.702 | 0.495 | 1.417 | 0.177 |
|  | HER2 Positive | -1.084 | 0.742 | -1.461 | 0.165 |
|  | ER Positive | 0.943 | 0.851 | 1.107 | 0.286 |
|  | PR Positive | -0.960 | 0.867 | -1.107 | 0.286 |
| **ZNF750** | Intercept | 5.979 | 1.441 | 4.148 | 0.0009 |
|  | Age | -0.018 | 0.011 | -1.565 | 0.139 |
|  | BMI | 0.017 | 0.029 | 0.577 | 0.573 |
|  | Tumor Grade | 0.466 | 0.292 | 1.599 | 0.131 |
|  | Cancer Stage | 0.345 | 0.185 | 1.864 | 0.082 |
|  | HER2 Positive | -0.566 | 0.277 | -2.038 | 0.060 |
|  | ER Positive | 1.050 | 0.318 | 3.298 | 0.005 |
|  | PR Positive | -0.506 | 0.324 | -1.560 | 0.140 |

**Figure 11.**
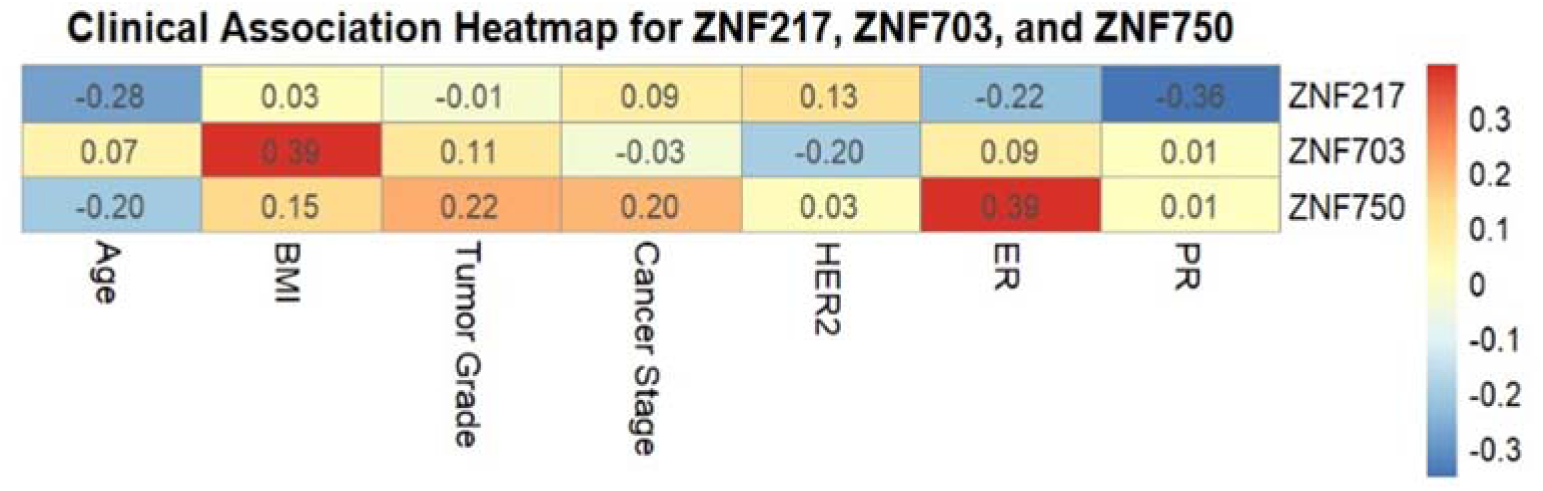
Clinical association heatmap of ZNF217, ZNF703, and ZNF750 expression. Heatmap showing correlation coefficients between tumor expression of ZNF217, ZNF703, and ZNF750 and clinical variables, including age, body mass index (BMI), tumor grade, cancer stage, HER2 status, estrogen receptor (ER) status, and progesterone receptor (PR) status. Color scale represents the strength and direction of correlations.

## DISCUSSION

This study provides new insights into the mutational and transcriptional landscape of ZNF217, ZNF703, and ZNF750 in breast cancer within a Kenyan cohort, addressing a key gap highlighted in previous research regarding underrepresented populations. The findings demonstrate that all three genes are both altered at the genomic level and transcriptionally dysregulated, supporting their proposed roles in breast cancer pathogenesis while also revealing important contrasts with existing literature.

The mutation landscape observed across ZNF217, ZNF703, and ZNF750 highlights gene-specific differences in both mutational burden and functional relevance. ZNF217 exhibited the highest mutation burden and a predominance of missense variants, including recurrent coding alterations and a truncating frameshift mutation, supporting its established role as an oncogenic regulator implicated in transcriptional dysregulation and tumor progression (Krig et al., 2010; Ansari-Pour et al., 2021). The spatial distribution of mutations across the protein, including within annotated zinc finger domains, further suggests potential structural and functional consequences. Notably, mutation frequencies in ZNF217 were consistent with TCGA data, reinforcing its relevance across independent cohorts.

In contrast, ZNF703 demonstrated a comparatively low mutation burden and limited recurrence, with variants distributed across the protein without clear clustering. The predominance of missense and synonymous mutations, coupled with minimal high-impact alterations, is consistent with previous studies indicating that ZNF703 contributes to tumorigenesis primarily through transcriptional activation and overexpression rather than frequent coding mutations (Klæstad et al., 2021; Zhang et al., 2022). The concordance of low mutation frequency between this cohort and TCGA further supports this interpretation.

ZNF750 displayed a high number of mutations within this cohort; however, these were largely composed of synonymous and low-impact variants, with most mutations occurring outside annotated functional domains. This pattern contrasts with reports describing ZNF750 as a tumor suppressor frequently inactivated by disruptive mutations (Butera et al., 2020; Cassandri et al., 2020). The absence of ZNF750 mutations in the TCGA dataset further suggests that the observed mutation profile may be cohort-specific. Collectively, these findings indicate that, while ZNF750 is frequently altered at the nucleotide level in this dataset, the predominance of low-impact variants and lack of domain enrichment point toward a more limited direct effect on protein function, potentially implicating alternative regulatory mechanisms.

The gene expression analysis further strengthens the evidence for the involvement of these genes in breast cancer. All three genes were significantly upregulated in tumor tissues, supporting previous studies that have linked overexpression of ZNF217 and ZNF703 to tumor aggressiveness and poor clinical outcomes (Krig et al., 2010; Marzbany et al., 2019). The consistent upregulation of ZNF217 observed in this study aligns with its role as a transcriptional regulator of cell survival and proliferation, including its involvement in activating ErbB3 signaling pathways (Krig et al., 2010). Similarly, the strong upregulation and high variability of ZNF703 expression are consistent with its known role in driving oncogenic signaling and tumor heterogeneity, particularly in luminal breast cancer subtypes (Klæstad et al., 2021; Zhang et al., 2022).

However, the expression pattern of ZNF750 contrasts with its traditionally described tumor-suppressive role. While previous literature suggests that loss or dysregulation of ZNF750 contributes to cancer progression through impaired differentiation (Butera et al., 2020; Cassandri et al., 2020), this study found consistent upregulation of ZNF750 in tumor tissues. This discrepancy suggests that ZNF750 may not function solely as a classical tumor suppressor and may instead have context-dependent roles in breast cancer. The coexistence of both hyper-expression and mutation in ZNF750 observed in this study supports earlier suggestions that its dysregulation can occur in multiple forms (Cassandri et al., 2020), but extends this understanding by demonstrating that upregulation may also be a prominent feature in certain populations.

The analysis of gene expression in relation to clinical variables revealed that most associations were weak, indicating that dysregulation of these genes is largely independent of traditional clinicopathological parameters. This finding suggests that ZNF217, ZNF703, and ZNF750 may act as intrinsic regulators of tumor biology rather than markers of disease stage or severity. However, two significant associations were identified: ZNF703 expression was positively associated with BMI, and ZNF750 expression was higher in ER-positive tumors. While previous literature has primarily focused on the oncogenic roles of ZNF703 and its association with aggressive tumor phenotypes (Marzbany et al., 2019), the observed link with BMI introduces a potential connection between metabolic factors and gene regulation. Similarly, the association between ZNF750 and ER status suggests a possible interaction with hormone signaling pathways, extending current understanding of its role beyond differentiation processes.

Although stratification of gene expression by intrinsic breast cancer subtypes (e.g., luminal, HER2-enriched, and triple-negative) could provide additional biological insight, such analyses were not performed in this study due to the limited sample size within each subtype group. The small number of samples per category would reduce statistical power and increase the risk of spurious associations. Nevertheless, the observed association between ZNF750 expression and ER-positive status suggests a potential link with luminal subtypes, which warrants further investigation in larger, well-powered cohorts.

Despite these strengths, several limitations should be considered when interpreting the findings. The relatively small sample size may have limited the statistical power to detect subtle associations, particularly in the analysis of gene expression and clinical variables. In addition, the study focused on three selected genes, which, although biologically relevant, do not capture the full complexity of breast cancer genomics. The reliance on computational predictions for mutation impact and RNA-seq–based expression analysis may also introduce methodological biases, and the cross-sectional nature of the study limits causal inference. Furthermore, the findings are derived from a single population, which may affect generalizability to other populations with different genetic and environmental backgrounds.

These limitations highlight important areas for future research. Larger, multi-center studies incorporating diverse populations are needed to validate these findings and improve generalizability. Integrative multi-omics approaches, including epigenomic and proteomic analyses, would provide a more comprehensive understanding of the regulatory networks involving ZNF genes. Functional studies are particularly important to clarify the biological roles of ZNF217, ZNF703, and especially ZNF750, whose behavior in this study differs from traditional expectations. In addition, further investigation into the observed associations with BMI and ER status may provide new insights into the interplay between metabolic, hormonal, and transcriptional pathways in breast cancer.

In conclusion, this study confirms and extends existing literature by demonstrating that ZNF217 and ZNF703 function as key oncogenic regulators in breast cancer, while revealing a more nuanced and potentially context-dependent role for ZNF750. Importantly, the findings provide population-specific evidence from a Kenyan cohort, supporting the notion that genetic and transcriptional features of breast cancer may vary across populations. This underscores the need for continued research in underrepresented groups to improve the accuracy of molecular characterization and support the development of precision oncology approaches.

## ACKNOWLEDGMENT

We would like to thank the patients for their consent to provide samples and Aga Khan University Hospital (Nairobi) and AIC Kijabe Hospital (Kijabe) for granting access to patient samples.

## DATA ACCESSIBILITY

WES data is accessible at SRA database Accession number: PRJNA913947, while RNA-seq data is accessible at the GEO database under Accession number: GSE225846. All other datasets for this study are included in the article’s Supplementary Material.

## AUTHORS’ CONTRIBUTION

FM conceived the idea and designed the research project, collected the samples and assembled experiment materials. MK, GW, JG, and KM performed the data analysis. FM and GW wrote the manuscript. ARH, SS, SA helped with drafting and reviewing the manuscript. All authors contributed to the revision and final editing of the manuscript prior to submission.

## SUPPLEMENTARY MATERIALS

Table S1: ZNF mutation summary

Table S2: ZNF mutation positions

Table S3: Amino acid change summary

Table S4: TCGA mutation dataset

Table S5: ZNF protein domains

Table S6: Combined ZNF expression and pathological and clinical data

